# Morphological brain alterations and morphological brain network disorganizations in heroin and methamphetamine abstinent patients

**DOI:** 10.1101/2024.11.12.623173

**Authors:** Xiaoliang Zhou, Zhifu Zhang, Haitao Yuan, Yuzhou Wang, Ziyu Xu, Wenbin Liang, Mingwu Lou

## Abstract

Heroin and methamphetamine are the two common types of drugs abused, which poses significant health risks. However, the neural mechanisms underlying the effects of drug addiction on human brain are unclear. In this study, we collected T1-weighted magnetic resonance imaging data from 26 heroin abstinent (HA) patients, 24 methamphetamine abstinent (MA) patients and 32 healthy controls. Four surface-based morphological features including cortical thickness (CT), fractal dimension (FD), gyrification index (GI), and sulcal depth (SD) were calculated, and further used to construct the morphological brain networks. We observed the common CT reductions of the right TE 1.0 and TE 1.2 and SD reductions of the right intermediate lateral area 20 for HA and MA patients, HA-specific CT reductions in the left area 2, and the MA-specific GI reductions in the left medial area 6 and right dorsomedial parietooccipital sulcus. For the morphological brain networks, HA patients exhibited the global disorganizations (higher shortest path length) in CT-based networks, whereas MA patients showed the disrupted nodal efficiency of the left medial area 38 in CT-based networks, the right caudal area 7 in GI-based networks, and the right inferior occipital gyrus in SD-based networks. Furthermore, the altered SD of HA patients and disrupted nodal efficiency of MA patients were associated with drug abuse-related clinical variables. Our findings suggest the morphological index-dependent effects of drug addiction on human brain morphology, and indicate the differential neural mechanism underlying heroin and methamphetamine abuses which attack the global and local information transfer of morphological brain networks, respectively.

## Introduction

Drug addiction poses significant health risks, including severe neurological and psychological damage, leading to the devastating consequences to individuals and to society (1, 2). In China, heroin and methamphetamine are the two most common types of drugs abused. Numerous studies have investigated the effects of these two drugs on human from various perspectives, including clinical characteristics, behavioral manifestations, neurocognition, and genetics (3–7), for promoting effective addiction treatment. In recent years, an increasing number of studies have focused on the neural mechanisms of heroin and methamphetamine acting on the human brain using neuroimaging techniques (8–10).

Structural magnetic resonance imaging (MRI) is a popular way to characterize the brain morphology, due to its low cost, high test-retest reliability, and high spatial resolution. Through this technique, studies show that drug addiction can lead to the brain morphological alterations in heroin abstinent (HA) and methamphetamine abstinent (MA) patients (11–13). For example, reduced gray matter density of the right middle frontal gyrus is found in the both short-term and long-term MA patients. Meanwhile, less gray matter density reduction is observed in the long-term MA patients than short-term MA patients, which is correlated with executive function (14). It should be noted that, most of these studies only focus on single morphological feature, like gray matter density or cortical thickness (CT). In addition to these two features, many other morphological features offer nonredundant descriptions of brain morphological characteristics that exhibit differential developmental and aging trajectories and are differently affected in various brain diseases (15–17). Therefore, examining the effects of drug addiction on multiple brain morphological features in HA and MA patients can help to deepen understand the underlying neural mechanism.

Recently, structural MRI has been proved to contain abundant brain connectivity information, i.e., morphological similarity (18, 19). To date, many approaches have been proposed to estimate the statistical interdependence of local morphological features between regions (19). The resultant morphological brain networks have become a growing focus of brain network studies. Among these approaches, the divergence-based individual morphological brain network has been widely used for studying the coordinated patterns of brain morphology in health and disease, due to its high test-retest reliability, phenotypic relationship, and neurobiological substrates (20, 21). Therefore, from the perspective of divergence-based individual morphological brain network, exploring the human brain network disorganizations in HA and MA patients can bring a new light into the neural mechanism underlying drug addiction.

In this study, we aimed to explore the effects of drug addiction on brain morphology and morphological brain networks in HA and MA patients. We first examined the morphological alterations at the regional level among HA patients, MA patients and healthy controls (HCs), using four surface-based features including CT, fractal dimension (FD), gyrification index (GI), and sulcal depth (SD). Subsequently, the morphological brain networks were constructed based on these four features, respectively, to examine the global and local disorganizations of HA and MA patients. Finally, the patient-related morphological alterations and network disorganizations were used to correlate with clinical variables.

## Methods

### Participants

A total of 26 HA and 24 MA patients were recruited from a drug rehabilitation hospital in Shen Zhen, China. Thirty-two HCs were enrolled from the local community and matched by age, education level, and handedness. This study followed the principles of the Declaration of Helsinki and was approved by the Ethics Committee of Longgang Central Hospital. All subjects were informed of the procedure and provided informed consent before the start of the study.

The inclusion criteria for patients with HA and MA included the following items: (a) diagnosed with a substance use disorder according to the criteria in the Diagnostic and Statistical Manual of Mental Disorders, 4^th^ edition (DSM-IV); (b) abstinent for less than six months; (c) no substance-use history except for heroin, methamphetamine, nicotine, and alcohol; (d) right-handed; and (e) 18 to 45 years old. The recruitment criteria for the HCs included the following: (1) good health with no severe diseases; (2) no history of substance abuse other than nicotine and alcohol; (3) right-handed; and (4) 18 to 45 years old. The exclusion criteria for all participants were as follows: (a) a history of mental disorders; (b) severe injury by collision, such as head trauma; (c) current medical illness; or (d) claustrophobia or other contraindications for magnetic resonance imaging (MRI).

The demographic information (age and gender) was collected for each participant, and the clinical information (duration of drug abuse, frequency of drug abuse, duration of abstinence, dosage of drug use per time, dosage of drug use per day, cigarettes per day, anxiety score, and depression score) was collected for each patient.

### Image acquisition

All data were acquired on a 3 T Siemens Prisma with a 64-channel head coil. The T1-weighted images were obtained with the following parameters: repetition time = 2300 ms, echo time = 2.32 ms, field of view = 240 × 240 mm^2^, matrix = 256 × 256, slice thickness = 0.45 mm, 208 sagittal slices, interslice gap = 0.45 mm, voxel size = 0.9 × 0.9 × 0.9 mm^3^, and flip angle = 120°. During the scanning, the subjects were instructed to lie down immobile, wear noise-cancelling earbuds, and close the eyes.

### Structural image preprocessing

All structural images were preprocessed using the standard pipeline with the CAT12 toolbox (http://dbm.neuro.uni-jena.de/cat12/) based on the SPM12 package (https://www.fil.ion.ucl.ac.uk/spm/software/spm12/). The CAT12 toolbox was demonstrated to offer a fast and reliable alternative to the FreeSurfer when performing the analysis of surface-based morphometry (22). First, each structural image was segmented into gray matter, white matter, and cerebrospinal fluid based on an adaptive Maximum A Posterior technique (23), after spatial adaptive non-local means denoising filter and bias-correction for reducing the effects of potential artifacts. Then, CT was estimated using a projection-based thickness method (24), and FD, GI, and SD were calculated as the slope of a logarithmic plot of surface area versus the maximum l-value (25), absolute mean curvature (26), and Euclidean distance between the central surface and its convex hull, respectively, based on the spherical harmonic reconstructions. Finally, the individual CT, FD, GI, and SD maps were resampled into the common fsaverage template and smoothed using a Gaussian kernel (15-mm full width at half maximum for CT and 25-mm full width at half maximum for the others).

### Construction of morphological brain networks

Based on the vertex-wise morphological maps, single-subject morphological brain networks were constructed based on CT, FD, GI, and SD, respectively, by estimating morphological similarity in terms of distributions of morphological index.

#### Network nodes

To define the node of morphological brain network, a prior brain atlas was used to divide the cortex into a set of 210 regions of interest (ROIs) (27), with each ROI denoting a node.

##### Network edges

To define the edge of morphological brain network, Jensen-Shannon divergence (JSD)-based morphological similarity was used according to previous studies (20, 21). Specifically, for each individual morphological map, the values of all vertices within each ROI were extracted to estimate a probability density function using ksdensity function in MATLAB. Then, the resultant probability density functions were transformed to probability distribution functions (PDFs) for calculating the JSD between any pair of ROIs. Formally, for two PDFs *P* and *Q*, the JSD was calculated as:

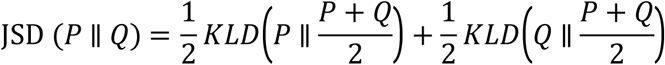

where 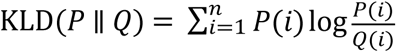 with *n* being the number of sample points which was set as 2^8^ according to previous studies (20, 21). After the transformation to a distance metric (square root) and subtraction from 1, the JSD represented the degree of similarity of morphological distributions between two ROIs (0, absolutely different; 1, exactly the same).

Finally, we obtained four 210 × 210 single-subject morphological index-dependent brain networks, i.e., CT-, FD-, GI-, and SD-based networks (CTN, FDN, GIN, and SDN) for each participant.

### Topological analysis of morphological brain networks

#### Thresholding

Before topologically characterizing the morphological brain networks, a thresholding procedure was used to convert them into binary networks, for excluding the effects of spurious connections on subsequent network analyses. Specifically, the sparsity-based threshold was utilized, which was defined as the ratio of the actual edge number to the maximum possible edge number. At a fixed sparsity, the sparsity-based thresholding procedure ensures the same number of connections across participants by applying a subject-specific threshold to exclude weak connections. Given the lack of a definitive standard for choosing a single sparsity, a consecutive sparsity range of [0.03 0.3] with an interval of 0.05 was used to guarantee that the resultant networks had sparse network properties (28, 29). Furthermore, to ensure that the resultant networks had no isolated nodes or multiple connected components, a minimum spanning tree algorithm was integrated into the thresholding procedure (30).

#### Network measures

For each morphological brain network derived above (82 participants × 4 morphological indices × 6 sparsity levels), we calculated the small-world coefficients, i.e., clustering coefficient (*Cp*) and characteristic path length (*Lp*), and nodal efficiency (*Ei*). Specifically, the small-world coefficients were further normalized (*zCp* and *zLp*) by the corresponding average value of 100 matched random networks, which were generated with a topological rewiring algorithm to preserve the same degree distribution as the actual networks (31). All the topological analyses were performed with the GRETNA toolbox (32).

### Statistical Analysis

#### Demographic variables

Age was examined using one-way analysis of variance with significance level estimated by permutation procedure (1,000 times), and gender was examined using chi-squared test.

#### Regional morphology

One-way analysis of variance was used to examine the differences of regional mean CT/FD/GI/SD among the three group. Specifically, the significance level was estimated by permutation procedure (1,000 times). To correct for multiple comparisons, a false discovery rate (FDR) procedure was used across 210 ROIs for each morphological index at the level of *q* < 0.05. For significant group main effects, post hoc tests were further performed with two-sample *T* tests followed by multiple comparison correction with the FDR procedure.

#### Network organization

One-way analysis of variance was used to examine the differences in topological organizations (*Cp*, *Lp*, *zCp*, *zLp*, and *Ei*) of morphological brain networks among the three group. Specifically, the significance level was estimated by permutation procedure (1,000 times). To correct for multiple comparisons, a false discovery rate (FDR) procedure was used across 210 ROIs for *Ei* in each morphological index at the level of *q* < 0.05. For significant group main effects, post hoc tests were further performed with two-sample *T* tests followed by multiple comparison correction with the FDR procedure.

#### Clinical relationship

To examine the relationships between clinical variables and regional morphology and network organization, Spearman rank correlation was used. Specifically, only the regional morphology and network organization showing significant differences in patient group compared to the HCs was used. The FDR procedure was used to correct for multiple comparisons in each patient group and each clinical variable.

## Results

### Demographic Characteristics

There were no significant differences in gender or age among the three groups (Table 1).

**Table 1.** Demographic and clinical variables.

| | HA<br>(n = 26) | MA<br>(n = 24) | HCS<br>(n = 32) | Group effect<br>*F* or $\chi^2$ (*P*) |
| --- | --- | --- | --- | --- |
| Gender (F/M) | 2/24 | 5/19 | 3/29 | 2.402 (0.301) |
| Age (years) | 31.13±6.47 | 33.04±6.98 | 30.39±7.94 | 1.844 (0.168) |
| Duration of abstinence (days)<sup>a</sup> | 140.26±246.65 | 112.75±155.09 | - | - |
| Duration of drug abuse (years)<sup>b</sup> | 6.95±4.41 | 7.39±6.69 | - | - |
| Frequency of drug abuse (times/day)<sup>c</sup> | 1.96±1.07 | 7.92±12.85 | - | - |
| Dosage of drug use per time (gram)<sup>c</sup> | 0.32±0.29 | 0.20±0.22 | - | - |
| Dosage of drug use per day (gram)<sup>c</sup> | 0.65±0.75 | 0.48±0.31 | - | - |
| Cigarettes per day (number)<sup>c</sup> | 17.00±9.55 | 11.17±6.70 | - | - |
| Anxiety score<sup>d</sup> | 7.17±6.10 | 8.28±9.25 | - | - |
| Depression score<sup>e</sup> | 5.76±4.51 | 7.00±5.81 | - | - |

Data are presented as mean ± standard deviation. HA, heroin abstinent; MA, methamphetamine abstinent; HCs, healthy controls; F, female; M, male.
<sup>a</sup>Data are missing for 7 HA patients.
<sup>b</sup>Data are missing for 4 HA patients and 1 MA patient.
<sup>c</sup>Data are missing for 3 HA patients.
<sup>d</sup>Data are missing for 8 HA patients and 6 MA patients.
<sup>e</sup>Data are missing for 8 HA patients and 3 MA patients.

### Abnormal morphology in the patients

#### CT

The left area 2 (*F* = 7.045, *P* < 0.001) and right TE 1.0 and TE 1.2 (*F* = 8.655, *P* < 0.001) were found to exhibit significant group main effects of regional mean CT among three groups. Post hoc tests revealed that CT of the left area 2 was lower in HA patients than MA patients (*T* = 3.590, *P* = 0.003) and HCs (*T* = 2.788, *P* = 0.008), and CT of the right TE 1.0 and TE 1.2 was lower in HA (*T* = 3.545, *P* < 0.001) and MA (*T* = 3.511, *P* < 0.001) patients than HCs (Fig 1).

**Fig 1.**
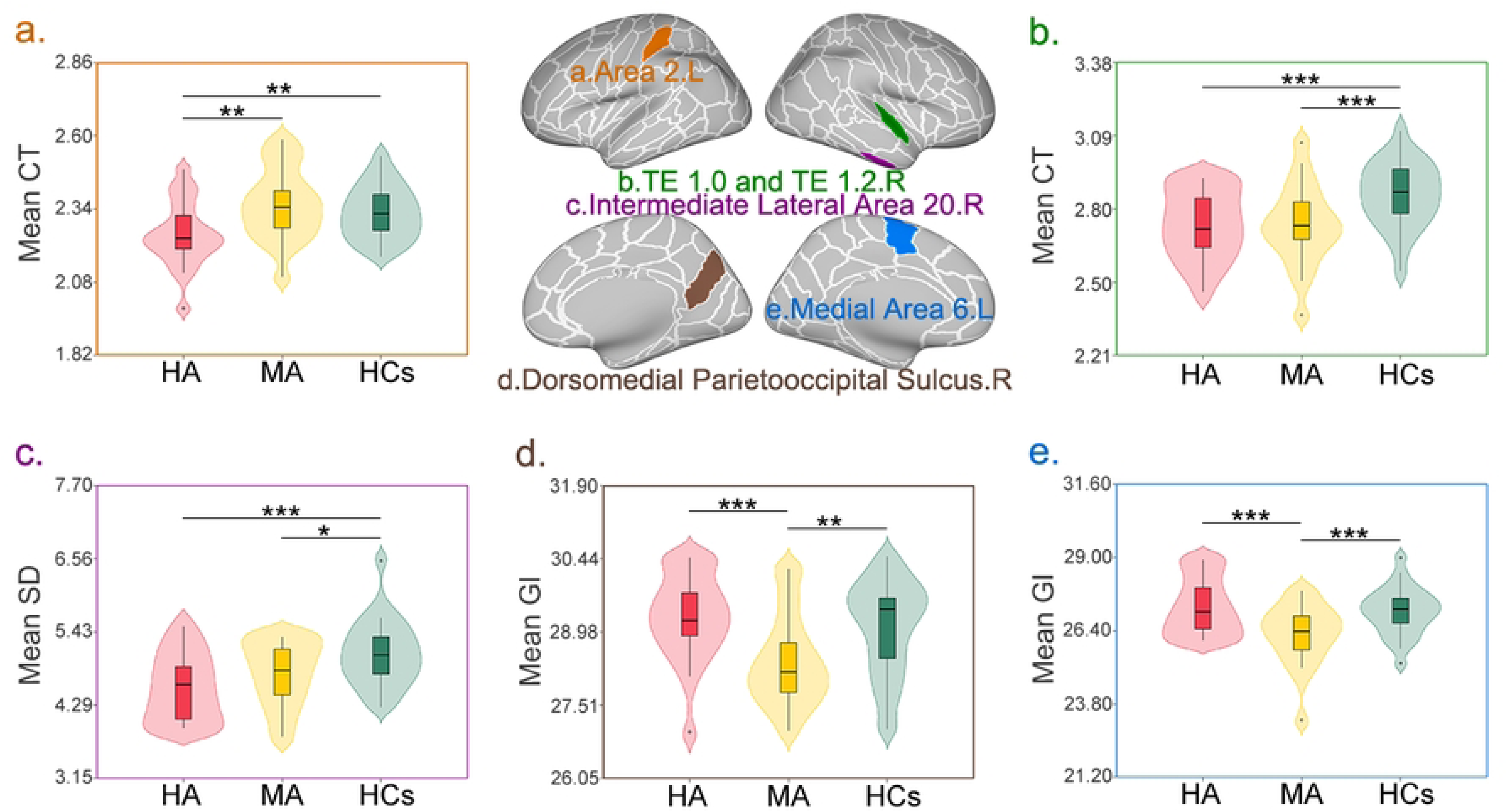
Abnormal cortical morphology in the patient groups. The common CT reductions of the right TE 1.0 and TE 1.2, and SD reductions of the right intermediate lateral area 20 for HA and MA patients, HA-specific CT reductions in the left area 2, and the MA-specific GI reductions in the left medial area 6 and right dorsomedial parietooccipital sulcus were observed. HA, heroin abstinent; MA, methamphetamine abstinent; HCs, healthy controls; CT, cortical thickness; SD, sulcal depth; GI, gyrification index; L, left; R, right.

#### FD

No significant group main effects of regional mean FD among three groups were observed in any region.

#### GI

The left medial area 6 (*F* = 8.256, *P* < 0.001) and right dorsomedial parietooccipital sulcus (*F* = 7.728, *P* < 0.001) were found to exhibit significant group main effects of regional mean GI among three groups. Post hoc tests revealed that GI of these two regions was lower in MA patients than HA patients (left medial area 6: *T* = 3.765, *P* < 0.001; right dorsomedial parietooccipital sulcus: *T* = 3.733, *P* < 0.001) and HCs (left medial area 6: *T* = 3.334, *P* < 0.001; right dorsomedial parietooccipital sulcus: *T* = 3.068, *P* = 0.009) (Fig 1).

#### SD

The right intermediate lateral area 20 was found to exhibit significant group main effects of regional mean SD among three groups. Post hoc tests revealed that SD of this region was lower in HA (*T* = 4.293, *P* < 0.001) and MA (*T* = 2.381, *P* = 0.020) patients than HCs (Fig 1).

### Global disorganizations of CTNs in the HA patients

Only the CTNs showed group main effects of *Lp* among three groups at the marginal significance level (*F* = 2.872, *P* = 0.058). Post hoc tests revealed that the *Lp* of CTNs was higher in HA patients than MA patients (*T* = 2.270, *P* = 0.044) and HCs (*T* = 1.826, *P* = 0.053) (Fig 2). No significant differences of small-world coefficients among three groups were observed in FDNs, GINs, or SDNs.

**Fig 2.**
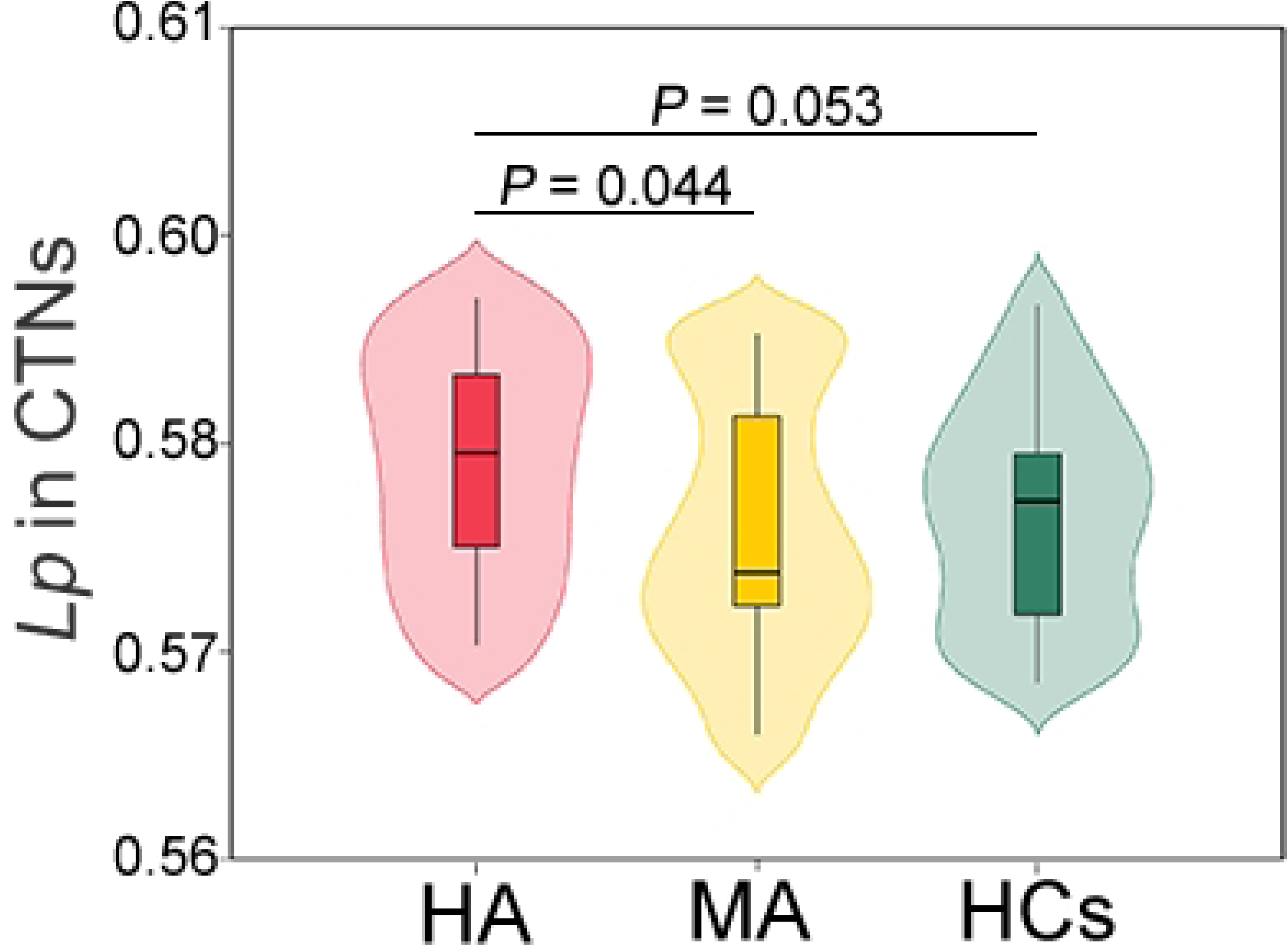
Global disorganizations of morphological brain networks in the HA patients. At the marginal significance level, HA patients exhibited the higher *Lp* of CTNs than MA patients and HCs. HA, heroin abstinent; MA, methamphetamine abstinent; HCs, healthy controls; *Lp*, shortest path length; CTNs, cortical thickness-based networks.

### Disrupted nodal efficiency of morphological brain networks in the MA patients

#### CTNs

The left medial area 38 was found to exhibit significant group main effects of *Ei* in the CTNs among three groups (*F* = 7.224, *P* < 0.001). Post hoc tests revealed that *Ei* was lower in MA patients than HA patients (*T* = 3.588, *P* = 0.002) and HCs (*T* = 3.007, *P* = 0.005) (Fig 3).

**Fig 3.**
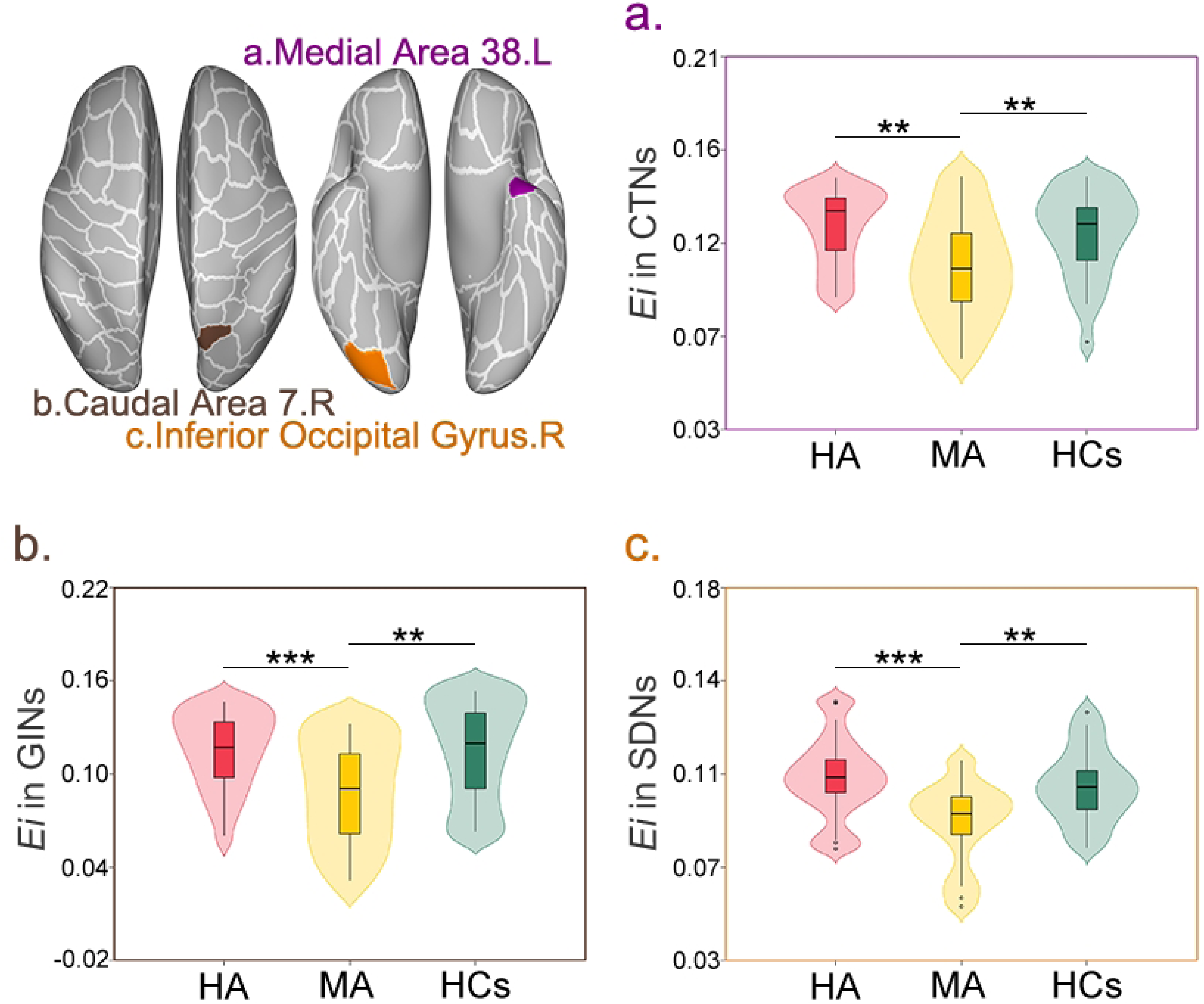
Disrupted nodal efficiency of morphological brain networks in the MA patients. MA patients showed the disrupted nodal efficiency of the left medial area 38 in CTNs, the right caudal area 7 in GINs, and the right inferior occipital gyrus in SDNs. HA, heroin abstinent; MA, methamphetamine abstinent; HCs, healthy controls; *Ei*, efficiency; CTNs, cortical thickness-based networks; GINs, gyrification index-based networks; SDNs, sulcal depth-based networks; L, left; R, right.

#### FDNs

No significant group main effects of *Ei* in the FDNs among three groups were observed in any region.

#### GINs

The right caudal area 7 was found to exhibit significant group main effects of *Ei* in the GINs among three groups (*F* = 8.310, *P* < 0.001). Post hoc tests revealed that *Ei* was lower in MA patients than HA patients (*T* = 3.486, *P* < 0.001) and HCs (*T* = 3.672, *P* = 0.003) (Fig 3).

#### SDNs

The right inferior occipital gyrus was found to exhibit significant group main effects of *Ei* in the SDNs among three groups (*F* = 10.517, *P* < 0.001). Post hoc tests revealed that *Ei* was lower in MA patients than HA patients (*T* = 4.304, *P* < 0.001) and HCs (*T* = 3.674, *P* = 0.003) (Fig 3).

### Clinical relationships of abnormal morphology and network disorganizations

#### HA patients

In the HA patients, we observed the significantly positive correlations between frequency of drug abuse and regional mean SD of the right intermediate lateral area 20 (*rho* = 0.560, *P* = 0.006). At the marginall significance level, the anxiety scores were negatively correlated with regional mean SD of the right intermediate lateral area 20 in the HA patients (*rho* = -0.530, *P* = 0.024) (Fig 4).

**Fig 4.**
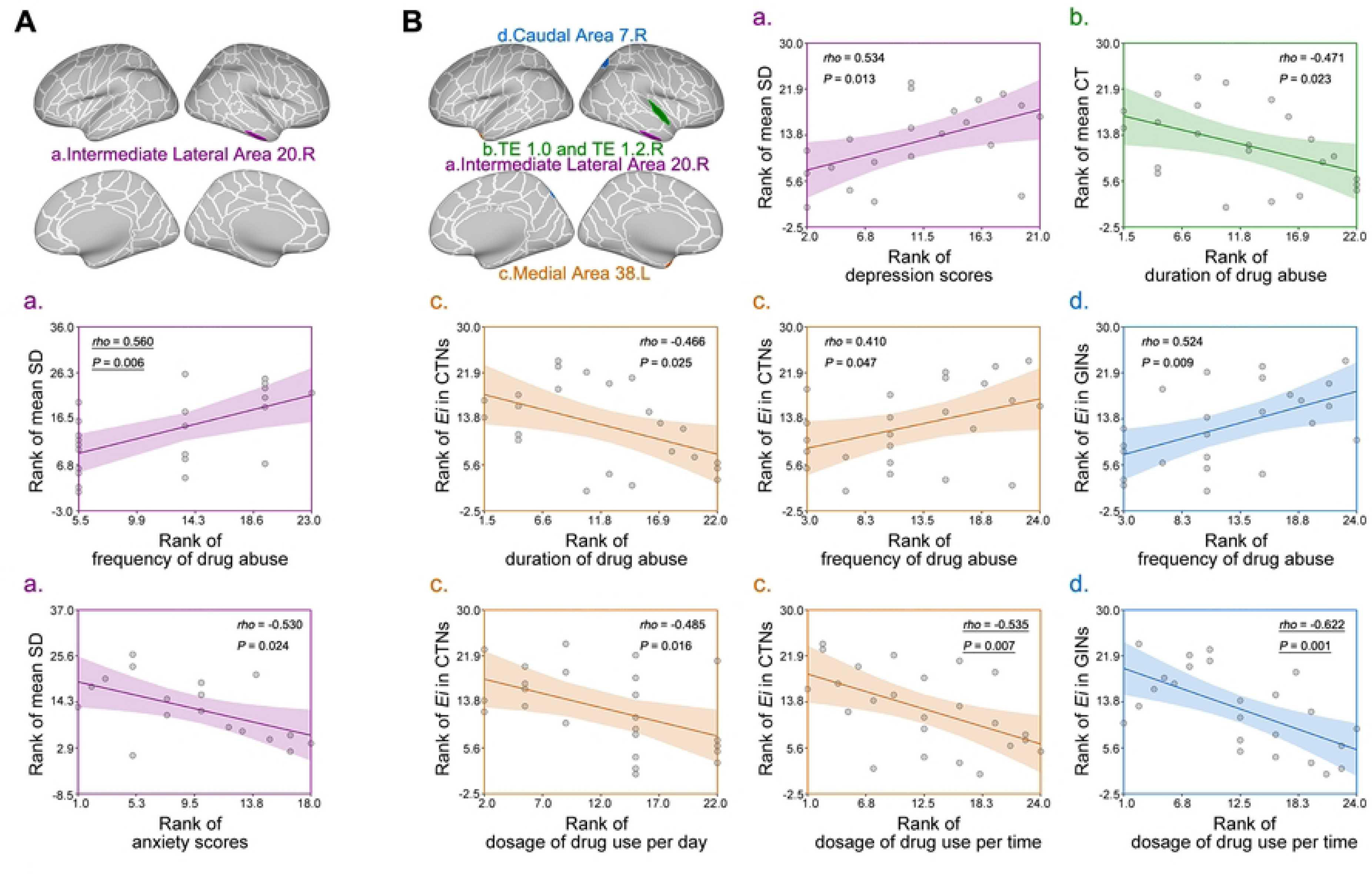
Clinical relationships of abnormal cortical morphology and network disorganizations in the heroin abstinent (**A**) and methamphetamine abstinent patients (**B**). CT, cortical thickness; SD, sulcal depth; *Ei*, efficiency; CTNs, cortical thickness-based networks; GINs, gyrification index-based networks; L, left; R, right. The *rho* and *P* with underline indicated the correlations with false discovery rate-corrected *P* < 0.05.

#### MA patients

In the MA patients, we observed the significant correlations of dosage of drug use per time with *Ei* of the left medial area 38 in CTNs (*rho* = -0.535, *P* = 0.007), and *Ei* of the right caudal area 7 in GINs (*rho* = -0.622, *P* = 0.001). At the marginall significance level, correlations of frequency of drug abuse with *Ei* of the left medial area 38 in CTNs (*rho* = 0.410, *P* = 0.047) and *Ei* of the right caudal area 7 in GINs (*rho* = 0.524, *P* = 0.009), correlations of duration of drug abuse with regional mean CT of the right TE 1.0 and TE 1.2 (*rho* = -0.471, *P* = 0.023) and *Ei* of the left medial area 38 in CTNs (*rho* = -0.466, *P* = 0.025), correlations of dosage of drug use per day with *Ei* of the left medial area 38 in CTNs (*rho* = -0.485, *P* = 0.016), and correlations of depression scores with regional mean SD of the right intermediate lateral area 20 (*rho* = 0.534, *P* = 0.013) were observed in the MA patients (Fig 4).

## Discussion

In this study, we characterized brain morphology by using multiple cortical-based morphological indices and constructing individual morphological brain networks. We observed the abnormal CT and SD in both HA and MA patients, and altered GI in MA patients. Additionally, HA patients exhibited the global disorganizations in CTNs, whereas MA patients showed local disrupted nodal efficiency in the CTNs, GINs, and SDNs. Furthermore, the altered SD of HA patients and disrupted nodal efficiency of MA patients were associated with drug abuse-related clinical variables. Our findings suggest the morphological index-dependent effects of drug addiction on human brain in HA and MA patients. Meanwhile, for the morphological brain networks, heroin influences the global organization, whereas methamphetamine attacks the local architecture.

In this study, we found that both the HA and MA patients exhibited abnormal CT and SD, and the MA patients showed altered GI. These findings suggest that the effects of drug addiction on human brain morphology in HA and MA patients are dependent on the morphological indices. The CT measures the thickness of cortex, and the GI and SD are the measures of cortical folding. These morphological indices offer nonredundant descriptions of cortical characteristics that exhibit differential developmental and aging trajectories and are differently affected in various brain diseases (15–17, 33). Such differences among the morphological indices may contribute to the observed morphological index-dependent effects of drug addiction in this study, which bring a new light into the neural mechanism underlying heroin and methamphetamine. Specifically, both the HA and MA patients exhibited the reduced CT of the right TE 1.0 and TE 1.2, and reduced SD of the right intermediate lateral area 20. The right TE 1.0 and TE 1.2 is a subregion of the superior temporal gyrus, and is involved in pain, anxiety, audition, speech (27). The right intermediate lateral area 20 locates in the inferior temporal gyrus, and plays an important role in visual association processes, e.g., emotional meaning of colors and facial expression (34). Hence, the abnormal morphology of these two regions in HA and MA patients may be a possible neural basis underlying the disruptions of related functions after drug addictions. It should be noted that, we observed the correlations of the SD of the right intermediate lateral area 20 with anxiety scores and depression scores in the HA and MA patients, respectively. These differences between HA and MA patients may indicate the differential mechanisms between drug effects of heroin and methamphetamine, although with the common SD reductions in HA and MA patients. Furthermore, we observed the HA-specific CT reductions in the left area 2, and the MA-specific GI reductions in the left medial area 6 and right dorsomedial parietooccipital sulcus. Given that the left area 2 participates in executive functions while both the left medial area 6 and right dorsomedial parietooccipital sulcus are involved in the vision and motion, the patient-specific morphological alterations in these regions are aligned with the differential cognitive deficits between HA and MA patients (7, 35).

In this study, HA patients exhibited the global disorganizations of CTNs, with higher *Lp* compared to HCs. *Lp* is the minimum number of edges that must be traversed to go from one node to another, and measures the global efficiency of parallel information transfer (36). Hence, the observed higher *Lp* in the HA patients suggests that the heroin abuse destroys some edges that support the global information transfer, resulting in the lower global efficiency of CTNs. While for MA patients, disruptions of nodal efficiency in CTNs, GINs and SDNs were observed. Such differences between the HA and MA patients indicate the differential neural mechanism underlying heroin and methamphetamine abuses, which attack the global and local information transfer, respectively. Specifically, for the MA patients, the reduced *Ei* of the left medial area 38 in CTNs was observed, and was correlated with the drug abuse-related variables (dosage of drug use per day, frequency of drug abuse, duration of drug abuse and dosage of drug use per time). The left medial area 38 is a subregion of the superior temporal gyrus, and is involved in the emotional processing, especially the happiness processes (27, 34). A study finds that the use of methamphetamine can predict the less happiness in the next year (37). Therefore, the correlations between the reduced *Ei* of the left medial area 38 in CTNs with the drug abuse-related variables in the MA patients may explain such prediction. Additionally, the MA patients also exhibited the reduced *Ei* of the right caudal area 7 in GINs, which was correlated with the dosage of drug use per day and frequency of drug abuse. The right caudal area 7 locates in the superior parietal lobule, and is involved in attention, executive functions and visuospatial processing (27, 34). These functions, especially the visuospatial processing, are reported to be disrupted in the MA patients (38, 39). Hence, the observed reduction of *Ei* in the right caudal area 7 in GINs may be the neural basis underlying the cognitive deficits in the MA patients. However, the mechanism among the methamphetamine abuse, cognitive deficits and disrupted nodal efficiency in GINs remains unclear. Future studies can investigate such mechanism to determine whether methamphetamine abuse causes the cognitive deficits via local disorganizations of GINs, or whether preexisting deficits of related cognition and disrupted nodal efficiency of GINs are susceptible to methamphetamine abuse, or both (39).

This study had several limitations. First, most participants in this study were male, so it is important to interpret the observed results cautiously, considering the potential gender differences in the effects of drug addiction on human brain, especially the methamphetamine (40–42). Thus, the findings of this study may not be fully generalizable to females, and future research should aim to include a more balanced sample with respect to the gender, for better understanding the effects of drug addiction on brain morphology. Second, no significant alterations in patient groups were observed in FD and FDNs. Such insensitivity of FD and FDNs to the effects of drug addiction warrants further investigation into the underlying mechanisms, e.g., distinct cellular mechanisms and genetic origins among the morphological indices and morphological brain networks (21, 43). Third, it should be noted that there is a lack of complete clinical data for some patients, which may reduce the statistical power of the observed clinical correlations, potentially affecting the reliability and generalizability of the results. Together with the relatively small sample size in this study, it is crucial to consider these limitations when interpreting our findings and to strive for more comprehensive data collection in future studies to strengthen the validity of clinical correlations.

## Acknowledgements

This work was supported by the National Natural Science Foundation of China (No. 81471661).

## Author contributions

Conceptualization, Xiaoliang Zhou, Wenbin Liang and Zhifu Zhang; Data curation, Xiaoliang Zhou and Wenbin Liang; Formal analysis, Ziyu Xu and Mingwu Lou; Funding acquisition, Mingwu Lou; Investigation, Mingwu Lou; Software, Xiaoliang Zhou, Wenbin Liang and Haitao Yuan; Supervision, Mingwu Lou; Writing – original draft, Xiaoliang Zhou and Wenbin Liang; Writing – review & editing, Xiaoliang Zhou, Yuzhou Wang, Ziyu Xu and Mingwu Lou.

## Data availability statement

All data that support the findings of this study are available from the corresponding author upon reasonable request.

